# Cellular chaining influences biofilm formation and structure in Group A *Streptococcus*

**DOI:** 10.1101/831123

**Authors:** Artur Matysik, Foo Kiong Ho, Alicia Qian Ler Tan, Anuradha Vajjala, Kimberly A. Kline

## Abstract

Group A *Streptococcal* (GAS) biofilm formation is an important pathological feature contributing to the antibiotic tolerance and progression of various GAS infections. Although a number of bacterial factors have been described to promote *in vitro* GAS biofilm formation, the relevance of *in vitro* biofilms to host-associated biofilms requires further understanding. In this study, we demonstrate how constituents of the host environment, such as lysozyme and NaCl, can modulate GAS bacterial chain length and, in turn, shape GAS biofilm morphology and structure. Disruption of GAS chains with lysozyme results in biofilms that are more stable. Based on confocal microscopy, we attribute the increase in biofilm stability to a dense and compact three-dimensional structure produced by de-chained cells. To show that changes in biofilm stability and structure are due to the shortening of bacterial chains and not specific to the activity of lysozyme, we demonstrate that augmented chaining induced by NaCl or deletion of the autolysin gene *mur1.2* produced defects in biofilm formation characterized by a loose biofilm architecture. We conclude that GAS biofilm formation can be directly influenced by host and environmental factors through the modulation of bacterial chain lengths, potentially contributing to persistence and colonization within the host. Further studies of *in vitro* biofilm models incorporating physiological constituents such as lysozyme may uncover new insights into the physiology of *in vivo* GAS biofilms.

## INTRODUCTION

*Streptococcus pyogenes*, or group A *Streptococcus* (GAS), is a human pathogen that can cause a wide variety of human diseases (1, 2). GAS infections and its associated complications, such as rheumatic heart disease and acute post-streptococcal glomerulonephritis, represent a significant public health burden and there is a continued interest in developing new treatment strategies against this pathogen (3, 4). Multiple lines of evidence show that biofilm formation during GAS infection is an important factor contributing to antibiotic treatment failure (5, 6). Furthermore, biofilm formation may play a role in GAS pathogenesis and disease by changing the bacterial transcriptional profile or by modulation of the host response (7-9). Investigations into GAS biofilm formation may yield new therapeutic targets and thus far, numerous bacterial factors, including virulence determinants such as M protein surface adhesin (10), pili (11, 12), capsule (10, 13), and secreted SpeB protease (14) have been implicated in GAS biofilm formation.

While chaining is a defining morphological feature of GAS, the contribution of GAS chaining to its pathogenicity and biofilm formation has not been deeply investigated. In other bacterial species, chaining contributes to virulence traits including complement evasion (15) and adhesion to host cells (16). In some instances, chaining has been found to contribute to biofilm formation. For example, the initial stages of *Bacillus subtilis* biofilm formation involves the formation of long chains of non-motile cells (17). For some strains of *Escherichia coli*, cellular chaining is important for structural maturation of biofilms and this is mediated by the autotransporter protein Ag43 (18). Interestingly, uropathogenic *E. coli* (UPEC) develop into biofilm-like intracellular bacterial communities (IBCs) in bladder umbrella cells and these bacteria subsequently emerge from host cells as long, filamentous forms (19, 20). In *E. faecalis*, deletion of the autolysin gene *atlA* results in an increase in chain length and a decrease in biofilm biomass (21-23). Attenuation in the virulence of *E. faecalis* Δ*atlA* in a zebrafish model of infection has been directly attributed to long-chain formation (24). Similarly, the autolysin LytB and its homologues have been identified in other streptococci, including *S. pneumoniae, S. gordonii* and *S. mutans*, as regulators of chain length, and deletion of the autolysin gene often decreased their capacity to form biofilms (25-27). Chaining induced by choline supplementation had been shown to inhibit *S. pneumoniae* biofilm formation, but the exclusive and mechanistic role of chaining in streptococcal biofilm formation has remained a topic for speculation (25).

It is poorly understood how GAS regulates its own chain length, but it is likely to be complex and highly dependent on environmental conditions. Growth medium supplemented with homologous antiserum containing anti-M antibodies induces the formation of long chains in GAS. Conversely, growth in medium supplemented with non-immune serum results in chain shortening (28), indicating that there are multiple components in serum that could differentially influence GAS chaining. A transposon library screen for genes influencing chain length in *S. sanguinis* found that a large number of transposon mutants, corresponding to 15% of the genes within the genome, had either a significant increase or decrease in bacterial chain length (29), suggesting that streptococcal chaining could be a highly regulated process. In GAS, the autolysin Mur1.2 is predicted to be involved in cell separation and morphology, due to its homology with LytB in *S. pneumoniae* (30). Transcriptomic studies showed that *mur1.2* is induced by Ser/Thr kinase Stk, an important regulator of cell division, growth and virulence (30). Furthermore, deletion of a transcriptional regulator *rgg2* resulted in overexpression of *mur1.2* and increased biofilm formation (31). We therefore hypothesized that the regulation of GAS chain length, either by environmental or genetic factors, plays a role in modulating biofilm formation and virulence.

In this study, we determined the contribution of chaining to GAS biofilm formation. We show that supplementation of growth medium with physiologically-relevant constituents, such as lysozyme or serum, can significantly shorten GAS chain length. Decreased GAS chaining subsequently alters biofilm morphology, producing a biofilm that is denser and more stable *in vitro*. By contrast, we show that augmented chaining, induced by either NaCl or deletion of growth phase-regulated autolysin gene *mur1.2*, results in biofilm that is more fragile and loose in structure. We show that the degree of chaining impacts GAS biofilm formation, that chaining can be differentially affected by factors in the local host environment, thereby not only demonstrating how host factors such as lysozyme can shape GAS biofilm formation *in vivo*, but also arguing for their consideration when developing physiologically relevant *in vitro* models of GAS biofilm formation.

## RESULTS

### Serum and lysozyme reduce chain length in GAS

We confirmed the previously reported de-chaining effect of non-immune serum by microscopic observation of multiple GAS strains (JS95, HSC5, JRS4, MGAS5005) cultured in serum-supplemented medium. For all strains, we observed a marked decrease in bacterial chain lengths when grown overnight in THY-G medium supplemented with 10% fetal bovine serum (FBS) as compared to growth in non-supplemented medium (**Figure 1a**). As lysozyme had been shown to abrogate *Bacillus cereus* chaining (32), we postulated that lysozyme could be a serum component that contributes to the shortening of GAS chains. Consistent with this idea, upon supplementation of growth media with 10 µg/ml of lysozyme, which is similar to concentrations found in human serum (33), we observed shorter GAS chains as compared to growth in non-supplemented medium (**Figure 1a**). At the same time, lysozyme did not significantly reduce GAS viability, as verified by Live/Dead staining (**Figure S1**), growth kinetics (**Figure S2a**), and CFU enumeration of cultures grown overnight in the presence of increasing lysozyme concentrations up to 1 mg/mL (**Figure S2b**). In conclusion, exposure to physiological levels of lysozyme reduces GAS chaining *in vitro* without an effect on GAS viability.

**Figure 1.**
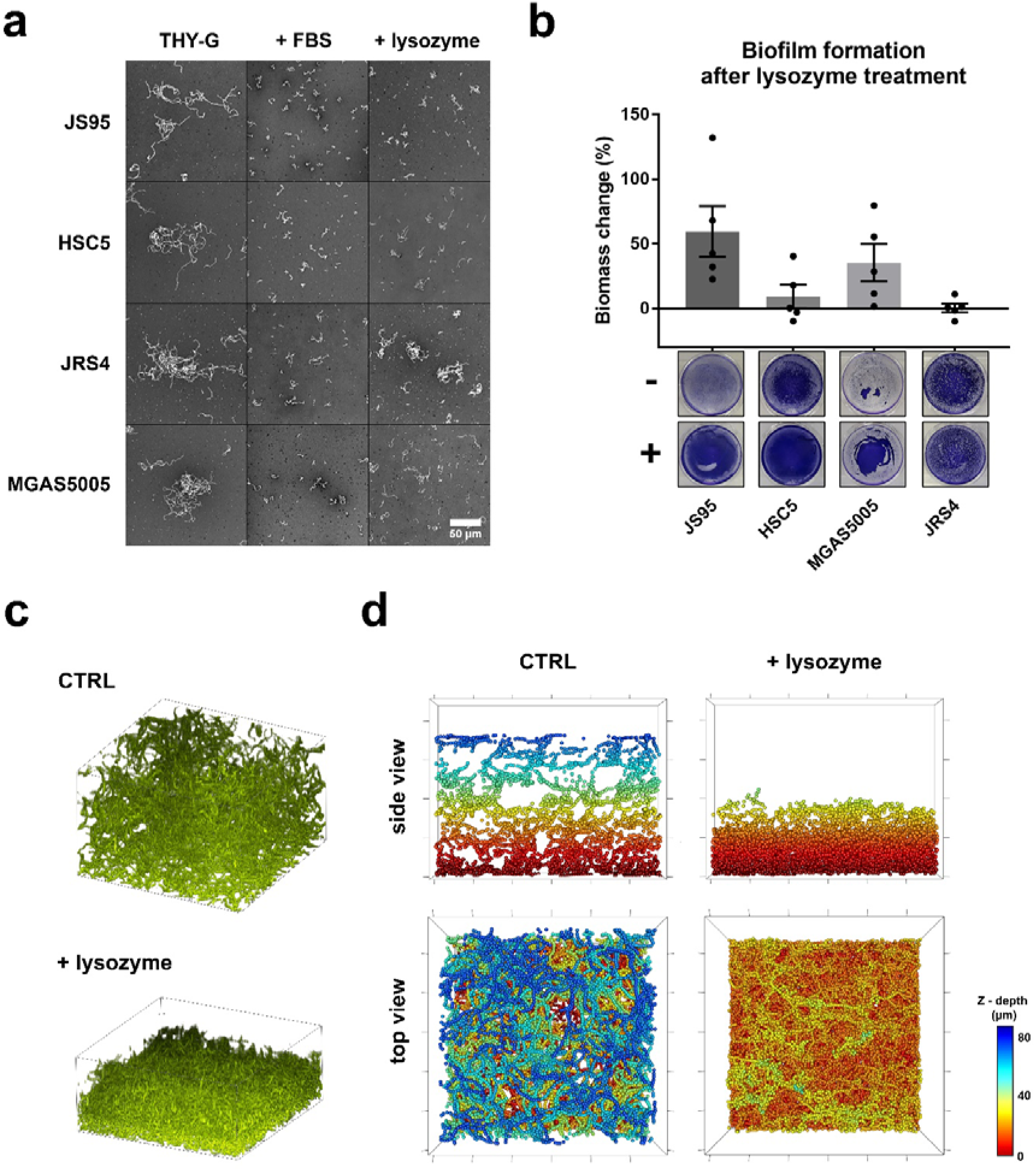
Lysozyme alters chaining and biofilm formation. **(a)** Overnight cultures of GAS strains JS95, HSC5, JRS4 and MGAS5005 in THY-G medium alone, supplemented with 10% FBS or lysozyme (10 µg/mL). Negative staining with nigrosin was used to visualize chaining and to minimize handling effects on the fragile chains. **(b)** CV staining and relative change in biofilm biomass after lysozyme supplementation.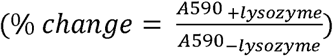. Biofilms were washed at high speed (250 rpm). Dots and error bars represent biological replicates and mean ± SEM, respectively. **(c)** Volume projection of CLSM stack and **(d)** Structure visualization of JS95 biofilm grown without and with 10 µg/mL lysozyme (coloured by Z-depth). Field of view (XY): 130 × 130 µm (volume projection and top view) and 130 × 10 µm (side view). Tick size: 20 µm.

### Reduction of chain length by lysozyme alters biofilm morphology

To investigate whether lysozyme-induced de-chaining influences biofilm formation, we performed crystal violet (CV) assays in 24-well polystyrene plates comparing static biofilms grown in the presence and absence of lysozyme. Preliminary CV assay experiments showed that JS95 biofilms grown in the presence of lysozyme had a more homogenous appearance compared to biofilms grown in the absence of lysozyme. As we thought that the change in biofilm morphology could represent a change in biofilm stability or structure, we modified the CV assay by the introduction of a vigorous washing step, which consisted of an additional 10 min washing on a rocker set to high speed (250 rpm). Biofilms of JS95, and to a smaller extent, MGAS5005 and HSC5, were more stable in the presence of vigorous washing (**Figure 1b**), as compared to control biofilms grown in medium without lysozyme supplementation. Although we did not observe changes for strain JRS4, the biofilm of JRS4 is relatively strong and dense even prior to lysozyme treatment, most likely due to combined effects of strong primary adhesion, tendency to form microcolonies (34), and low levels of capsule production which has been shown to improve static biofilm formation (13, 35, 36). Consequently, the JRS4 biofilm likely represents the upper limit of biofilm formation, which lysozyme is unable to further augment. Differences in biofilm stability were not detectable with washing at a lower speed (200 rpm, **Figure S2c**), indicating that lysozyme contributes to biofilm stability under high shear stress.

For a more in-depth investigation of the effect of lysozyme-induced de-chaining on biofilm morphology, we performed confocal microscopy and constructed three-dimensional representative views of GAS biofilms (**Figure 1c-d, S3**). Growth of JS95, HSC5 and MGAS5005 biofilms in the presence of lysozyme resulted in structural alterations characterized by tight and dense packing without obvious chaining. By contrast, in the absence of lysozyme supplementation, biofilms appeared as loosely packed bacterial chains forming thick, but not dense, biofilm structures. No obvious differences were noted for JRS4, which formed dense and coherent biofilms even prior to lysozyme treatment, as described above. Changes in biofilm ultrastructure correlate with the results obtained from CV assays performed under vigorous washing, providing a mechanistic explanation for how lysozyme could enhance GAS biofilm stability. GAS biofilms formed in the presence of lysozyme are made up of de-chained cells that can form more compact and stable biofilms, resulting in decreased susceptibility towards disruption by physical forces. Additionally, using the MBEC biofilm eradication assay, we observed that lysozyme treated cultures formed biofilms that were slightly more resistant to penicillin, while the MIC was not affected by lysozyme treatment (**Figure S4**).

### NaCl induces chaining and perturbs biofilm formation

High concentrations of NaCl in the growth medium induces chaining of *Listeria monocytogenes* possibly by inactivation or lowering expression of autolysins involved in cell division (37). We therefore examined JS95 cultures grown in various NaCl concentrations to see whether a similar effect occurs in GAS. Strain JS95 was chosen because it had a lower tendency to aggregate and form chains (**Figure 1a**), which could otherwise mask augmented chaining phenotypes. Supplementation of growth medium with NaCl to a final concentration of 0.9%, which is equivalent to that of normal saline, resulted in increased chaining, which appeared more stiff and rigid, as compared to cultures grown in non-supplemented growth medium (**Figure 2a**). Supplementation of growth medium with NaCl reduced biofilm formation in a dose-dependent manner (**Figure 2b**), and lysozyme was able to shorten chain length (**Figure 2a**) and restore biofilm biomass to the level of non-supplemented THY-G even in the presence of high NaCl concentrations (**Figure 2c**). We confirmed that changes in biofilm formation in the presence of lysozyme and/or NaCl were not a consequence of differences in adhesion properties by performing adhesion assays, in which normalized cell cultures were incubated in 24-well polystyrene plates for 1 hour, followed by washing and staining with crystal violet. Although we observed that NaCl increased adhesion slightly, addition of lysozyme did not have a significant effect (**Figure 2d**). NaCl supplementation to a total concentration of 0.9% did not affect growth nor cell viability (**Figure S1, S2a**). We observed similar effects of lysozyme and NaCl on biofilm formation assessed using staining by the nucleic acid dye Syto9, excluding the possibility that lysozyme or NaCl causes differential retention of crystal violet dye (**Figure S5**).

**Figure 2.**
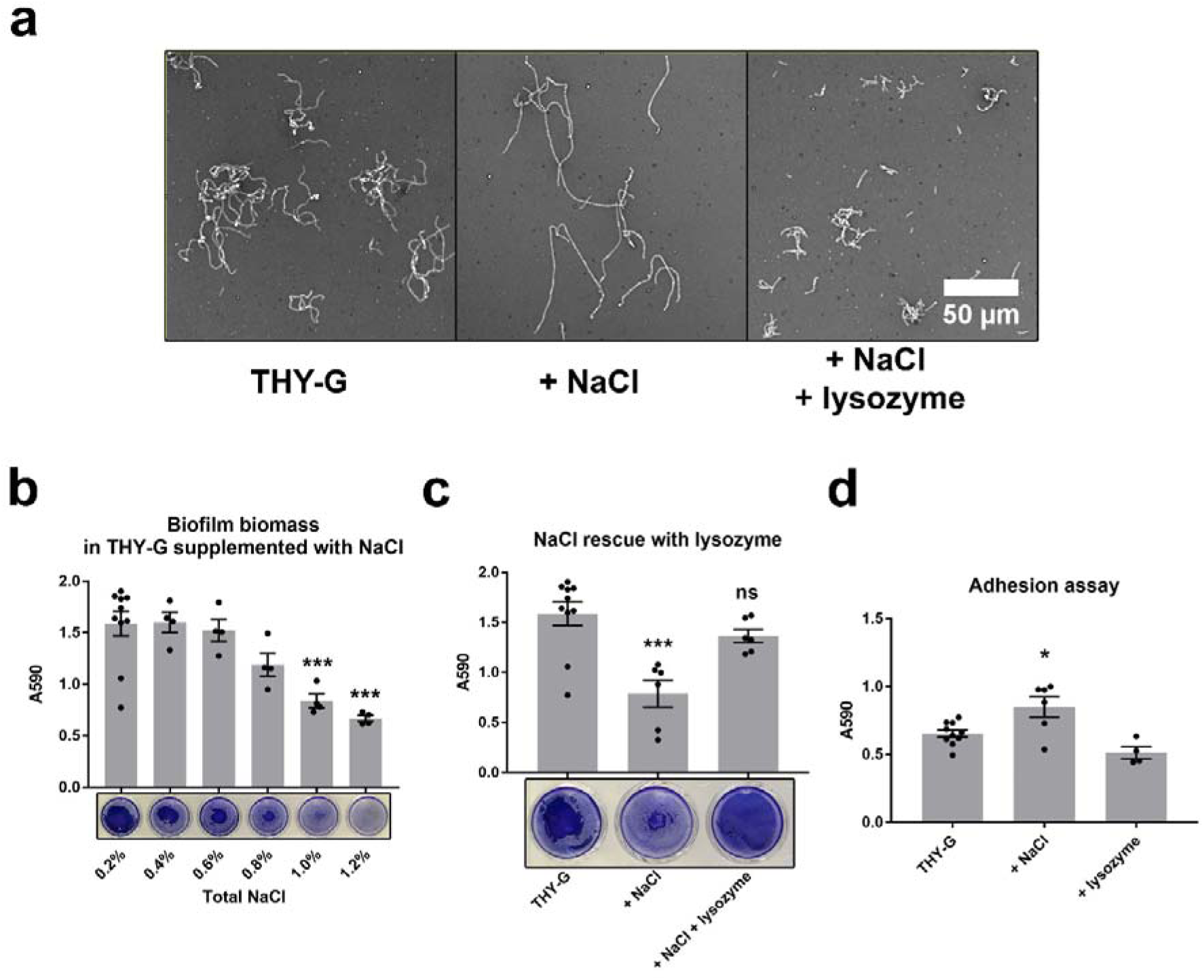
NaCl-induced chaining of GAS decreases biofilm formation. **(a)** Overnight cultures of GAS JS95 in THY-G medium alone, supplemented to a total of 0.9% NaCl, or supplemented with NaCl and 5 µg/mL lysozyme. **(b)** Biomass quantification and representative images of GAS JS95 biofilms grown in THY-G supplemented with increasing concentration of NaCl. **(c)** Biomass quantification and representative images of GAS JS95 grown in THY-G alone, supplemented with total 0.9% NaCl or supplemented with both 0.9% NaCl and lysozyme (10 µg/ml). **(d)** Adhesion assay of GAS JS95 grew in THY-G alone, supplemented with total 0.9% NaCl, or supplemented with lysozyme (10 µg/ml). Dots and error bars represent biological replicates and mean ± SEM, respectively, from at least three independent experiments. Statistical analysis was performed using one-way ANOVA, followed by Dunnett’s multiple comparisons test. *, *P* < 0.05; ***, *P* < 0.001; ns, non-significant.

### Autolysin Mur1.2 is involved in cell separation

In GAS, the gene *mur1.2* encodes a peptidoglycan hydrolase belonging to the glucosyl hydrolase family 73 (38), members of which have been experimentally demonstrated to exhibit N-acetylglucosaminidase activity (39). As the homologues LytB in both *S. pneumoniae* and *S. gordonii* (30, 40) are involved in cell separation and chaining, it had been predicted that Mur1.2 performs a similar role in GAS (30). To test this prediction, we constructed a non-polar in-frame deletion mutant of *mur1.2* in GAS strain JS95. Indeed, we observed that Δ*mur1.2* forms longer chains in planktonic culture as compared to WT (**Figure S6a**). The chain lengths of Δ*mur1.2* reverted to WT levels upon complementation, or could be further disrupted by lysozyme treatment (**Figure S6a, Figure 1a** and **Figure 2a**). We did not detect major differences in growth rates between WT and Δ*mur1.2* in THY medium (**Figure S6b**). Similarly, we did not detect any significant transcriptomic differences between WT and Δ*mur1.2* cultures grown to exponential phase by RNA-seq (**Figure S6c**), indicating that phenotypes associated with the deletion of *mur1.2* are not due to any off-target effects. Gene expression of *mur1.2* was, however, strongly regulated during growth in glucose-supplemented THY-G medium, with the peak of expression occurring in late exponential phase (**Figure S6d**).

### Mur1.2 alters biofilm phenotype

Scanning electron microscopy (SEM) revealed a long and rigid chain structure of the autolysin mutant biofilm (**Figure 3**). Similar to planktonic cultures, wild-type biofilm morphologies were restored in the *mur1.2* complemented strain. The lack of a visible matrix was consistent with our previous studies on the same strain, in which no matrix was detected by SEM as well as fluorescent imaging after wheat germ agglutinin (WGA) staining (13). Although we did not observe major defects in cell division even at high magnification using SEM (**Figure 3**), we noticed during SEM sample preparation that biofilms formed by Δ*mur1.2* were more fragile and easily disrupted by handling. Indeed, when assessed by CV biofilm assay, we found Δ*mur1.2* to be attenuated in biofilm formation as compared to WT. Similar to biofilms grown in 0.9% NaCl, biofilm formation of Δ*mur1.2* could also be restored to WT levels by lysozyme supplementation (**Figure 4**), suggesting that augmented chaining was directly responsible for the attenuation in the biofilm phenotype.

**Figure 3.**
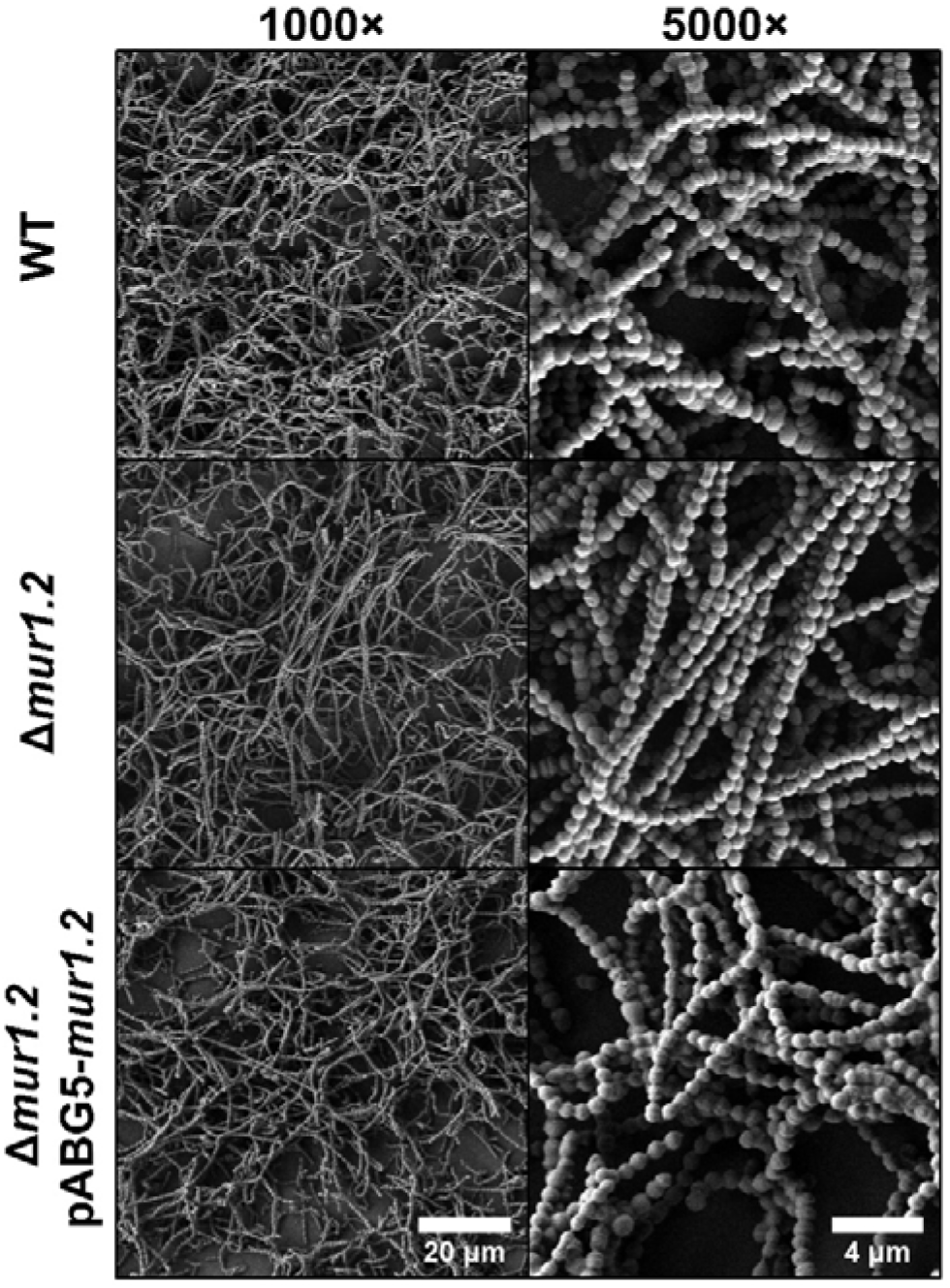
Deletion of *mur1.2* alters biofilm ultrastructure. Scanning electron micrographs (SEM) of JS95 WT, Δ*mur1.2*, and the *mur1.2* complemented strains biofilms grown on 6-well polystyrene plates in THY-G medium.

**Figure 4.**
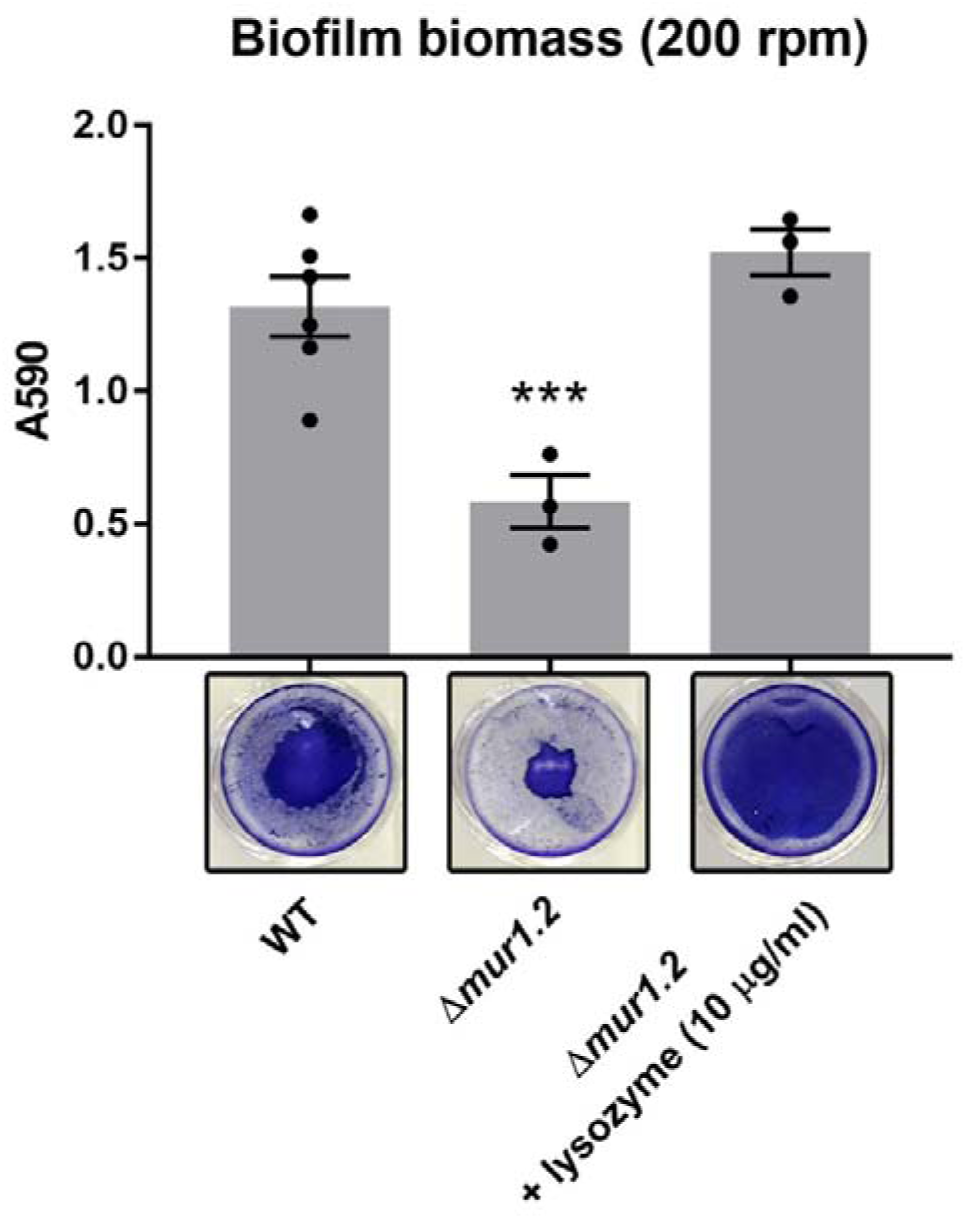
Attenuation in biofilm formation of *Δmur1.2* can be reversed by lysozyme treatment. GAS strains were grown for 24 hours before biofilm formation was quantified. Δ*mur1.2* biofilms were also grown in the presence of 10 µg/mL lysozyme to assess the ability to reverse attenuated phenotype. Biofilm formation was quantified using the CV assay, which was performed with an additional 10 min wash step with 1×PBS at 200 rpm. Dots and error bars represent biological replicates and mean ± SEM, respectively, from at least three independent experiments. Statistical analysis was performed using one-way ANOVA, followed by Dunnett’s multiple comparisons test. ***, *P* < 0.001.

### Mur1.2 deletion does not affect virulence in a NF *in vivo* model

Streptococcal species deficient in autolysins are often attenuated in virulence (41-44), which prompted us to investigate whether the same is true for GAS. To determine the importance of Mur1.2 in the context of invasive GAS infections, we used a mouse model of necrotizing fasciitis (NF) described previously (9), and chose strain JS95 as it was initially isolated from a NF patient (45). In this model, each flank of 3-4 week old BALB/c mice was injected subcutaneously with 10^8^ CFU of an early exponential GAS culture (WT, Δ*mur1.2*) or PBS as a control. However, we did not observe any differences in CFU at 24 hours post infection (**Figure 5a**). Immunostaining of lesion biopsy sections with anti-GAS antibody revealed the same spatial organization as described previously (9), with dense GAS aggregates located in the fascia (**Figure 5b**). However, we were not able to discern differences in chaining behaviour, and instead both mutant and wild-type bacteria appeared as cell aggregates. Taken together, the augmented chaining phenotype of Δ*mur1.2* grown *in vitro* in THY media is not apparent in a NF *in vivo* model, suggesting that de-chaining factors present *in vivo* may mask possible chain-associated phenotypes.

**Figure 5.**
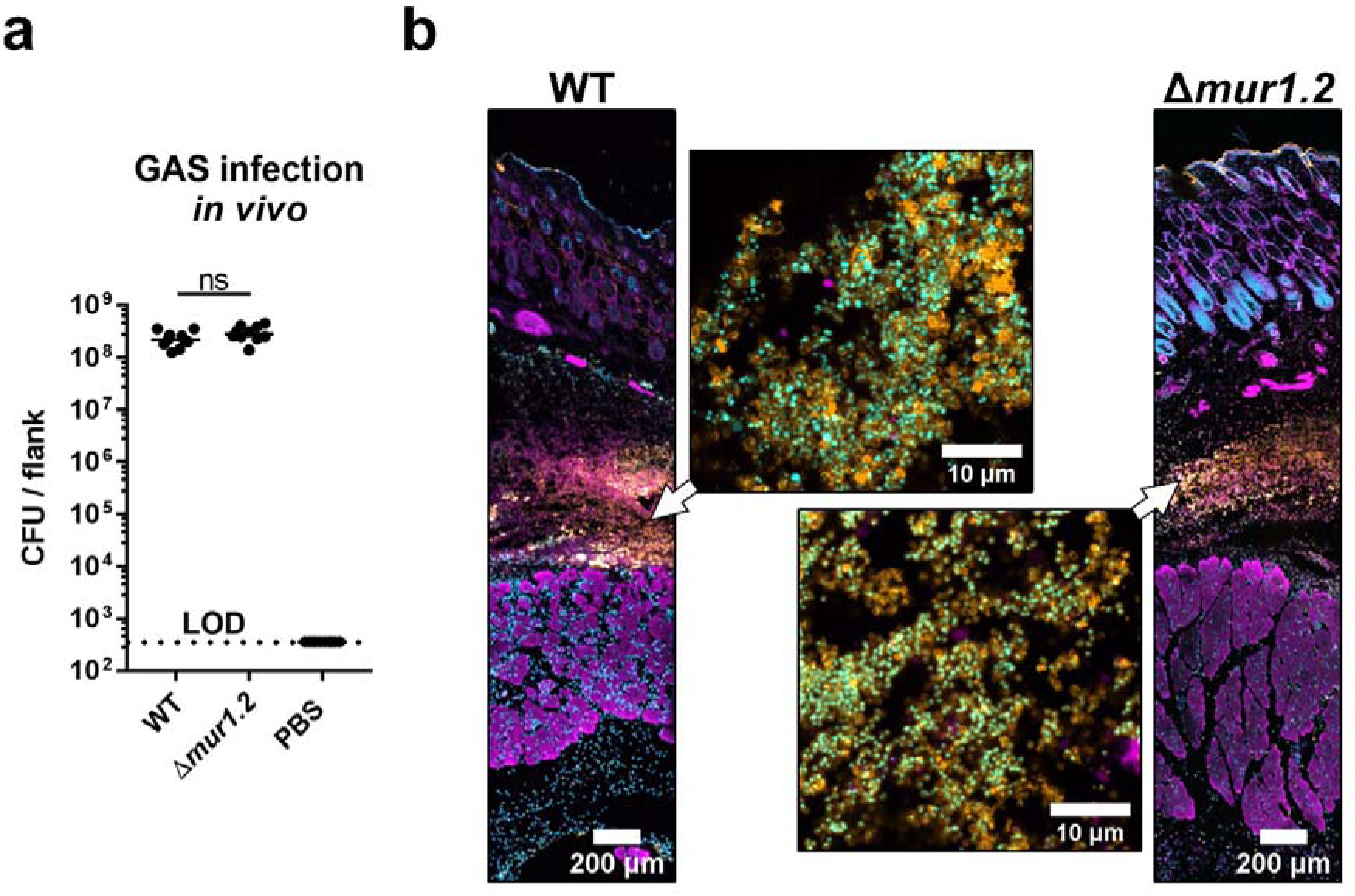
GAS *in vivo* infection. **(a)** BALB/c mice were infected subcutaneously with JS95 WT and JS95 Δ*mur1.2* and CFU recovered from lesions was quantified at 24 hours. n = 10 (WT, Δ*mur1.2*), n=4 (PBS), where n = number of flanks, 2 flanks / mouse. Comparisons between groups were performed using the Mann Whitney *U* Test. Dots and error bars represent biological replicates and median with interquartile range, respectively. **(b)** Biopsy sections stained with Hoechst33342 (DNA) – cyan, anti-GAS FITC (GAS) – orange, phalloidin-AF568 (actin) – magenta.

## DISCUSSION

### Chaining affects GAS biofilm formation and morphology

In this study, we confirmed previous reports that the addition of non-immune serum to growth medium shortens GAS chains (28). Similarly, lysozyme supplemented at levels typically observed in human serum resulted in the de-chaining of GAS cells. Thus, we concluded that lysozyme plays a major, but not necessarily exclusive, role in the de-chaining effect of non-immune serum. As lysozyme can act on bacteria as both a muramidase and as a cationic antimicrobial peptide (46), the precise mechanism by which lysozyme reduces GAS chain length is still not known. We discovered that lysozyme-induced de-chaining of GAS had a significant effect on static biofilm formation, as biofilms were more resistant to vigorous washes during CV biofilm assays. Careful analysis of biofilm structure by confocal microscopy revealed that bacterial cells within biofilms grown in the presence of lysozyme were dense and tightly packed. By comparison, GAS cultures grown in the absence of lysozyme produced biofilms with a loose and relaxed structure, also resulting in slightly reduced antibiotic tolerance. In other words, a thick *in vitro* biofilm phenotype due to chaining can carry a loose and fragile architecture, whereas a thin but dense and stable biofilm is composed of de-chained cells. An important implication of these observations is that apparent similarities in biofilm biomass as measured by standard assays may mask structural and functional differences in biofilm architecture.

Due to potential pleiotropic effects of lysozyme, we took an alternative approach and investigated how augmented chaining affects GAS biofilm formation. We found that supplementation of growth medium with NaCl not only augmented chaining of JS95, but also reduced the amount of static biofilm formation as compared to non-supplemented medium. Decreased biofilm formation was also observed for the hyper-chaining GAS JS95 mutant deficient in the autolysin Mur1.2. We showed that Mur1.2, although required for cell separation, is not crucial for cell division nor overall viability. Our observation that Δ*mur1.2* has decreased biofilm formation is consistent with the study of Zutkis et al., which showed both increased expression of *mur1.2* and biofilm formation upon mutation of the rgg2 transcriptional regulator (31). In both the NaCl-induced chaining and hyper-chaining mutant Δ*mur1.2*, biofilm formation could be restored by de-chaining with lysozyme. Although indirect effects of *mur1.2* deletion that are not associated with chain length might also affect biofilm formation, we were not able to identify any to date. RNA sequencing showed no transcriptomic effect of *mur1.2* deletion. SEM imaging, as well as fluorescent WGA staining of the same parent strain showed no visible matrix (13), suggesting that matrix production (e.g. eDNA release due to cell lysis) is not attenuated in the autolysin mutant. Moreover, unlike in other biofilm forming bacteria, direct evidence for eDNA in GAS biofilms has not been established (47). Further, we observed neither growth differences nor differences in viability between WT, Δ*mur1.2* mutant, and lysozyme treated WT that would suggest cell lysis.

Taken together, we showed that augmented chaining significantly changes biofilm architecture making it thick but at the same time loose and more fragile. Conversely, reduced chaining renders biofilms that are thinner but much more stable, due to tight cell packing.

### Host environment modulates chaining and biofilm structure

In this study, we were unable to confirm the role of chaining in virulence *in vivo* using a highly virulent NF-isolate GAS strain JS95, in a relevant well-established mice model of NF (9, 45). No differences in CFU were detected at 24 hours post infection. However, we observed that both WT and the hyper-chaining Δ*mur1.2* mutant did not produce obvious chaining phenotype in infected lesions. Although not tested directly in this study, the similar phenotype may be expected in pharyngeal colonization models previously developed for GAS (48). Several virulence factors have been identified as important for pharyngeal colonization and persistence, such as hyaluronic acid capsule (49) or SpeB (50), and most of them are under regulation of the CovRS two-component system (51). Despite CovRS controlling approximately 15% of the genome, its mutation was not found to directly affect *mur1.2* expression (30). Our studies showed no major transcriptomic effect of *mur1.2* deletion, suggesting altogether only a minor role of the Mur1.2 autolysin in the colonization. The lack of chaining of Δ*mur1.2 in vivo* suggests that the host environment has a dominant effect on GAS chaining. Indeed, GAS biofilms from *in vivo* studies appear as aggregates or clumps of bacterial cells with the absence of clear chaining (9, 10, 52, 53), morphologically distinct from *in vitro* biofilms grown without de-chaining agents, such as lysozyme or serum, as discussed above (5, 34, 36, 54). Notably, long GAS chains have been observed in macrophages after internalization and cytosolic replication (55), suggesting that internalization can protect GAS from de-chaining factors. Additionally, our results suggest that cytosolic concentrations of NaCl may be a contributing factor to long chain formation observed within host cells. It may be possible that the long chain phenotype can be found in non-phagocytic cell types as well and may be a feature of GAS intracellular infection. Although the role of long-chain formation in the GAS intracellular infection is not known, the GAS virulence factor SpeB is known to be regulated by NaCl, suggesting that GAS is able to sense and respond to NaCl during infection (56).

We propose that GAS biofilm formation during infection or colonization is heavily influenced by host factors such as lysozyme. Our results demonstrate that GAS biofilms grown in the presence of lysozyme were more structurally stable, robust, and tolerant to antibiotics, and suggest a mechanism by which GAS biofilms may become persistent *in vivo* and thereby render GAS infections recalcitrant to treatment. At the same time, the chaining phenotype and loose biofilm structure induced by NaCl or anti-M antibodies (28), autolysin down-regulation, or lack of de-chaining factors (e.g. lysozyme), might be desired for biofilm dispersal, bacterial spread, or colonization.

### Chaining behaviour is a physiologically-relevant aspect of GAS biofilm formation

Our results indicate that bacterial chaining is a caveat to the interpretation of *in vitro* biofilm assays for GAS, and possibly other streptococcal species, that could lead to missed or misinterpreted phenotypes. Autolytic activity has been positively correlated with the ability to form biofilms *in vitro* for *S. pneumoniae, S. gordonii* and *S. mutans* (25-27), but the direct role of chaining in inhibiting streptococcal biofilm formation has been unclear as some long-chaining strains also form biofilms (25). We showed that the effect of chaining on biofilm formation may not be easy to detect in the classical CV assays and is strongly dependent on handling (e.g. washing intensity) or assay setup (e.g. well size), due to abovementioned differences in biofilm density. In recent studies, GAS biofilm formation and host interactions were investigated using serum-containing cell culture medium, or in the presence of host cells (9, 57, 58). In support of developing physiologically-relevant GAS biofilm models, we demonstrated that biofilms grown in the presence of lysozyme were morphologically similar to GAS biofilms observed *in vivo*. Consequently, we propose that the inclusion of de-chaining factors, such as lysozyme or serum, in GAS growth medium should be considered as control conditions when designing GAS biofilm assays to better simulate physiological conditions.

## Conclusion

The results of this study provide insight and a note of caution for the *in vitro* investigation of *in vivo*-relevant GAS biofilm. We showed that GAS chaining significantly affects biofilm morphology and, through a modified CV assay, we demonstrated that GAS chaining also influences biofilm stability. Further investigation into these physiologically-relevant biofilms will not only inform how GAS causes persistent infections, but also accelerate the development of anti-biofilm therapies against GAS.

## MATHERIALS AND METHODS

### Bacterial strains and growth conditions

Bacterial strains and plasmids used in this study are listed in Table S1. E. coli strains were routinely grown in LB broth (BD Difco™) at 37°C with agitation and GAS strains were grown in Todd-Hewitt broth (Fluka) supplemented with 0.2% yeast extract (BD Bacto™) (THY) at 37°C with 5% CO_2_. For E. coli, erythromycin (Erm) was added at 500 µg/mL and kanamycin (Kan) at 50 µg/mL. For GAS, Erm was added at 1 µg/mL and Kan at 250 µg/mL.

### Construction of the Δ*mur1.2* deletion mutant

Chromosomal deletion of the *mur1.2* gene (BVA28_01865) from the JS95 genome (59) was performed by allelic replacement as previously described, using the temperature-sensitive shuttle vector pGCP213 (60). To create the deletion allele, 500 bp regions upstream and downstream of *mur1.2* were amplified using the primers listed in Table S2. The fragments were sewn together by overlap extension PCR and subsequently ligated into pGCP213 using the restriction sites BamHI and KpnI. After electroporation of the plasmid into competent GAS JS95 cells, successful transformants were selected with erythromycin at 30°C. Selection for chromosomal integration was performed at 39°C in the presence of erythromycin. For excision of the integrated plasmid, the cells were passaged without antibiotics once at 30°C, followed by several passages at 37°C. Mutants with the deletion were identified by colony PCR and confirmed by Sanger sequencing. The result is an in-frame deletion of most of the *mur1.2* gene, so that the deletion allele only contained the first two and last five codons.

### Complementation of the Δ*mur1.2* knockout strain

Complementation was performed by cloning the *mur1.2* gene, including its putative ribosomal binding site and transcription termination regions, into the E. coli – S. pyogenes shuttle vector pABG5 (61). A DNA fragment, starting from 21-bp upstream and ending at 130-bp downstream of the *mur1.2* gene, was amplified using primers containing arms homologous to pABG5 (Table S2). Cloning of the insert into linearized vector was performed using the In-Fusion HD Cloning Plus kit (Takara Bio), which placed the *mur1.2* gene under the control of the rofA promoter (61). The constructed plasmid was electroporated into competent cells of the Δ*mur1.2* mutant and selection of transformants was performed using kanamycin.

### RT-qPCR and RNA sequencing

RNA was extracted from planktonic cultures grown in THY-G at various stages of growth as previously described (13). RT-qPCR was performed using 2xSYBR FAST qPCR universal MasterMix kit (Kappa Biosystems, USA). Gyrase A (gyrA) was used as an endogenous control (62). The following primers were used (5’ > 3’): *mur1.2* (FWD) GGCTTATACGCTACAGACACCA; (REV) GTCATGCGTTTAGCCCAAGC; gyrA (FWD) CAACGCACGTAAGGAAGAAA; (REV) CGCTTGTCAAAACGACGTTA. For RNA sequencing, purified total RNA was depleted of ribosomal RNA using the Ribo-Zero™ rRNA removal kit (Illumina, USA). RNA and DNA concentrations were determined with the Qubit 2.0 fluorimeter (Invitrogen, USA), while RNA integrity was determined with the Agilent 2200 TapeStation (Agilent Technologies, Inc., USA). Only RNA with RIN values above 8.0 and DNA contamination of less than 10% were used for downstream analysis. Synthesis of cDNA was performed with the NEBNext RNA First-strand and NEBNext Ultra directional RNA Second-strand synthesis modules (New England BioLab, US). 250 bp paired reads were obtained on the HiSeq2500 system, and differential expression analysis was performed using edgeR package (63).

### Crystal violet (CV) biofilm assay

CV assay was performed as described in our previous studies (9, 13) with minor modifications. Briefly, growth medium (THY + 0.5% glucose, referred in text as THY-G) was inoculated 1:100 with overnight culture in THY, transferred to 24-well polystyrene plates and incubated statically for 24 hours at 37°C in a 5% CO_2_ atmosphere. Depending on the experiment, growth medium was supplemented with heat-inactivated fetal bovine serum (Gibco), chicken egg white lysozyme (Sigma) or sodium chloride (Sigma). Although fetal bovine serum was heat-inactivated at 56°C for 30 minutes, enzymatic activity of lysozyme was largely unaffected under these conditions **(Figure S7).** Concentrations of NaCl used in experiments are reported as weight by volume (w/v) and include NaCl already present in THY medium. Biofilms were then washed with PBS using a rocker set to either 200 or 250 rpm (as indicated in text), stained with 0.1% CV, washed again with PBS and dissolved in 96% ethanol. Biomass was quantified by absorbance measurement at 590 nm using the Infinite^®^ 200 Pro plate reader (TECAN). For biofilm assays, the eluent was diluted 5× in 96% ethanol to remain within the linear detection range of the spectrophotometer. As a second assay of biofilm formation, we modified the assay by substituting crystal violet staining with 500µl of 5µM Syto9 (Thermo-Fisher, USA), which after 10 min staining was similarly washed out and then resuspended thoroughly (to fully solubilize the biofilm) with 500µl PBS. Relative fluorescence was then measured (Ex 480 nm / Em 510 nm) using the Infinite^®^ 200 Pro plate reader (TECAN).

### Adhesion assays

Overnight cultures of GAS strains were inoculated, at a 1:100 dilution, into fresh THY-G medium containing either 0.9% total NaCl or 10 µg/mL lysozyme. After an overnight growth with either NaCl or lysozyme, overnight cultures were pelleted at 8,000 × g for 10 minutes and re-suspended in PBS, followed by normalization to an OD600 of 1.0. To assess for adhesion, 1 mL of normalized bacterial suspensions were dispensed into wells of a 24-well plate in triplicate and then incubated at 37°C for 1 hour. After incubation, wells were washed 3× with PBS and the remaining adherent bacteria was quantified by the CV assay.

### Growth kinetic and viability assays

For growth kinetic assays, overnight cultures of GAS strains were re-suspended in PBS, followed by inoculation into fresh THY medium at a 1:100 dilution and grown at 37°C with 5% CO_2_. At every hour for 12 hours, 1 mL of bacterial culture was transferred to a cuvette and absorbance was measured at 600 nm in a UV-VIS spectrophotometer (UV-1240, Shimadzu). For viability assays, overnight cultures of GAS strains were used to inoculate, at a 1:100 dilution, fresh THY medium containing varying concentrations of chicken egg white lysozyme. The prepared cultures were then incubated for 16 hours at 37°C with 5% CO_2_. Prior to plating on THY agar plates for CFU enumeration, bacterial chains were disrupted by centrifugation at 8,000 × g for 10 minutes.

### Imaging

Chaining of overnight planktonic cultures was captured by negative staining, in which 5 µL of 10% w/v nigrosin solution was mixed with 5 µL of bacterial culture, smeared on microscope glass slide, air-dried and images were acquired using transmitted light mode of the Leica SP8 microscope. Negative staining was used for better visualization of chaining and to minimize the effect of handling on bacterial chains. CLSM imaging of biofilms was performed as previously described (13). Briefly, biofilms in 8-well imaging chambers (Ibidi, Germany), were gently washed with PBS, fixed with 4% paraformaldehyde (PFA) for 15 min at room temperature, and stained with Syto9 (Thermo-Fisher, USA) according to manufacturer’s protocol. To preserve the spatial structure, biofilms were not left to dry at any step of the procedure. Z-stacks were collected with a Leica SP8 microscope equipped with 63×/NA1.4 oil immersion lens. Volume projections were rendered using DAIME software (64) and the morphology visualization was done using FIJI and R (65) as described before (66). Tissue biopsies were fixed in 4% PFA for 4 h at room temperature, incubated in 30% sucrose overnight at 4°C, frozen in OCT compound (Sakura, California, USA) using liquid nitrogen, cryosectioned to thickness of 15 µm using a Leica CM1860 cryostat (Leica Biosystems, Germany) and transferred to poly-L-lysine coated microscope slides. Sections were then stained with FITC-conjugated anti-GAS polyclonal antibody (1:20, cat#PA1-73060, Thermo-Fisher, USA), phalloidin-A568 (cat#A12380, Thermo-Fisher, USA) and Hoechst33342 (Thermo-Fisher, USA) according to manufacturer protocol, mounted using Vectashield medium (Vector Laboratories, USA) and imaged using a Zeiss LSM780 microscope (Carl Zeiss, USA) with 20×/NA0.80 and 100×/NA1.4 objectives for low and high magnification, respectively.

### Scanning Electron Microscopy

Biofilms grown in 6-well polystyrene plates were washed three times with 0.1 M PB, then fixed overnight with 2.5% glutaraldehyde in 4°C. 1% OsO_4_ was used for post-fixation, after which samples were dehydrated in an ethanol gradient, dried with HMDS (Sigma Aldrich), and coated with platinum. Imaging was conducted using Jeol 7610F (Jeol, USA).

### Mouse model of necrotizing fasciitis

The virulence of wild-type and Δ*mur1.2* strains was assessed using a mouse model of NF as previously described (9). Briefly, cultures were grown to early-log phase (OD = 0.2) in THY media, re-suspended in PBS and normalized. The flanks of 3-4 week old female BALB/c mice (InVivos, Singapore) were shaved and disinfected prior to subcutaneous injection of a 100 µL inoculum containing 1 × 10^8^ CFU. At 24 hours post infection, the resultant lesion was excised with an 8 mm biopsy punch (Robbins Instruments. Inc., New Jersey, USA). Biopsies were then homogenized in sterile PBS prior to CFU enumeration. All procedures were approved and performed in accordance with the Institutional Animal Care and Use Committee (IACUC) of Nanyang Technological University (NTU), School of Biological Sciences (ARF SBS/NIE [A0327]).

### Statistical data analysis

Statistical analyses were performed using the GraphPad Prism 6 software. Comparisons between two groups were performed using the two-tailed unpaired Student’s *t-*test. Comparisons between multiple groups were performed using ordinary one-way ANOVA with the Dunnett’s multiple comparison test. CFU enumeration data from mice infection experiments were analysed with the two-tailed Mann-Whitney *U* test. *P* values less than 0.05 were considered statistically significant.

## ACKNOWLEDGMENTS

We would like to thank Dr. Emanuel Hanski for providing bacterial strains used in this study and the members of the Kline lab for assistance with experiments. We express our gratitude to the SCELSE sequencing facility, especially Daniela Moses and her colleagues for library preparation and RNA sequencing. The authors acknowledge financial support from the Singapore Centre for Environmental Life Sciences Engineering (SCELSE), whose research is supported by the National Research Foundation Singapore, Ministry of Education, Nanyang Technological University and National University of Singapore, under its Research Centre of Excellence Programme. This work was also supported by the National Medical Research Council under its Clinical Basic Research Grant (NMRC/CBRG/0086/2015).

