## Supplementary figures and tables for "Cellular chaining influences biofilm formation and structure in Group A *Streptococcus*"

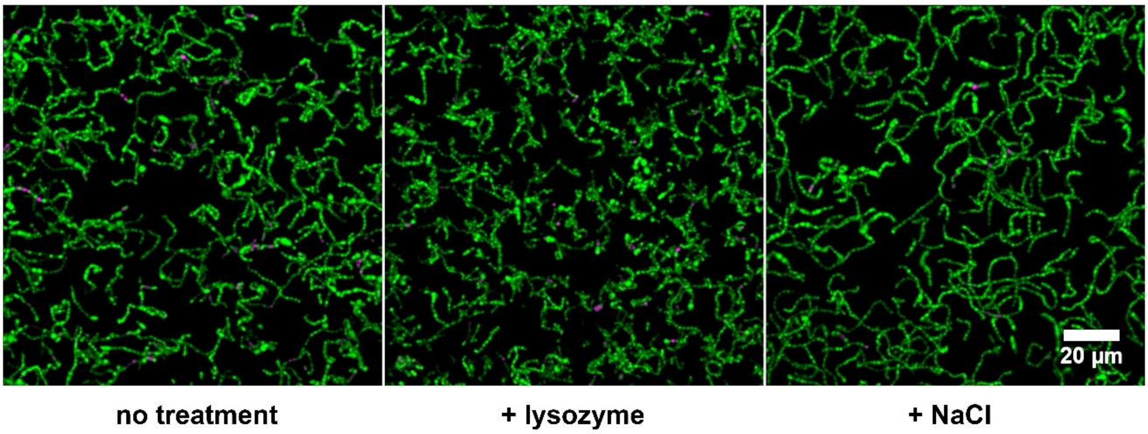


**Figure S1.** Live/Dead staining of planktonic cultures supplemented with NaCl (total 0.9%) or lysozyme (10 µg/ml). Pseudo-colouring with green and magenta represents live and dead cells, respectively.

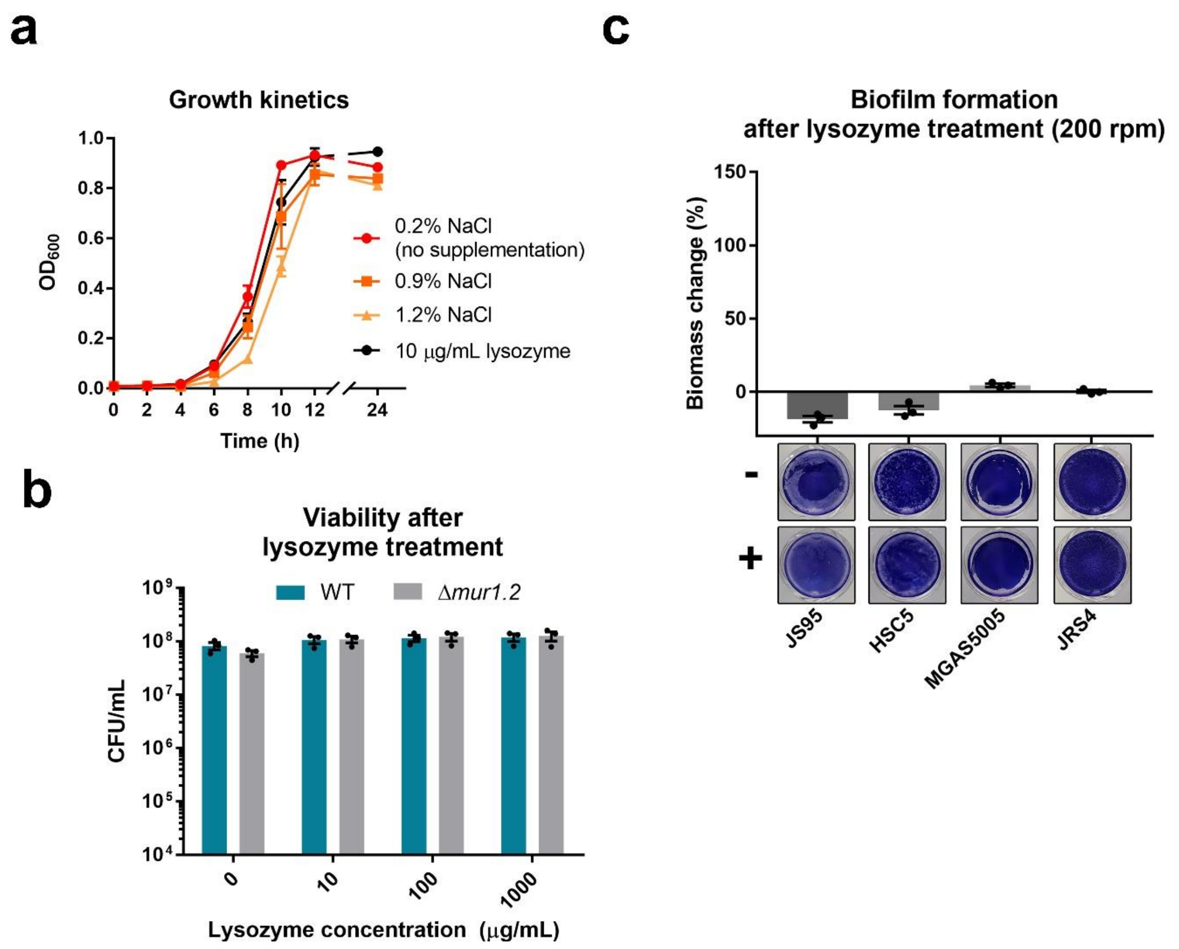


**Figure S2.** **(a)** Growth kinetics of planktonic cultures supplemented with NaCl (accounting for and including the 0.2% NaCl present in THY medium) or lysozyme (at 10 µg/ml). **(b)** CFU enumeration of JS95 WT and Δ*mur1.2* cultures after 16 hours of growth in THY medium containing increasing concentrations of lysozyme. Apparent, but not statistically significant, differences between CFU counts at 0 µg/mL lysozyme versus all other concentrations are most likely due to residual chaining, despite disruption of chains by centrifugation prior to plating. Dots and error bars represent biological replicates and mean ± SEM, respectively, from three independent experiments. Statistical analysis was performed using one-way ANOVA, followed by Dunnett's multiple comparisons test. **(c)** Changes in GAS biofilm formation after low speed wash. CV staining and relative change in biofilm biomass in THY-G medium with supplementation of 10 µg/mL lysozyme ($\%\ change = \frac{A590_{+lysozyme}}{A590_{-lysozyme}}$). Biofilms were washed at low speed (200 rpm) for 10 minutes on a rocker. Dots and error bars represent biological replicates and mean ± SEM, respectively, from three independent experiments.

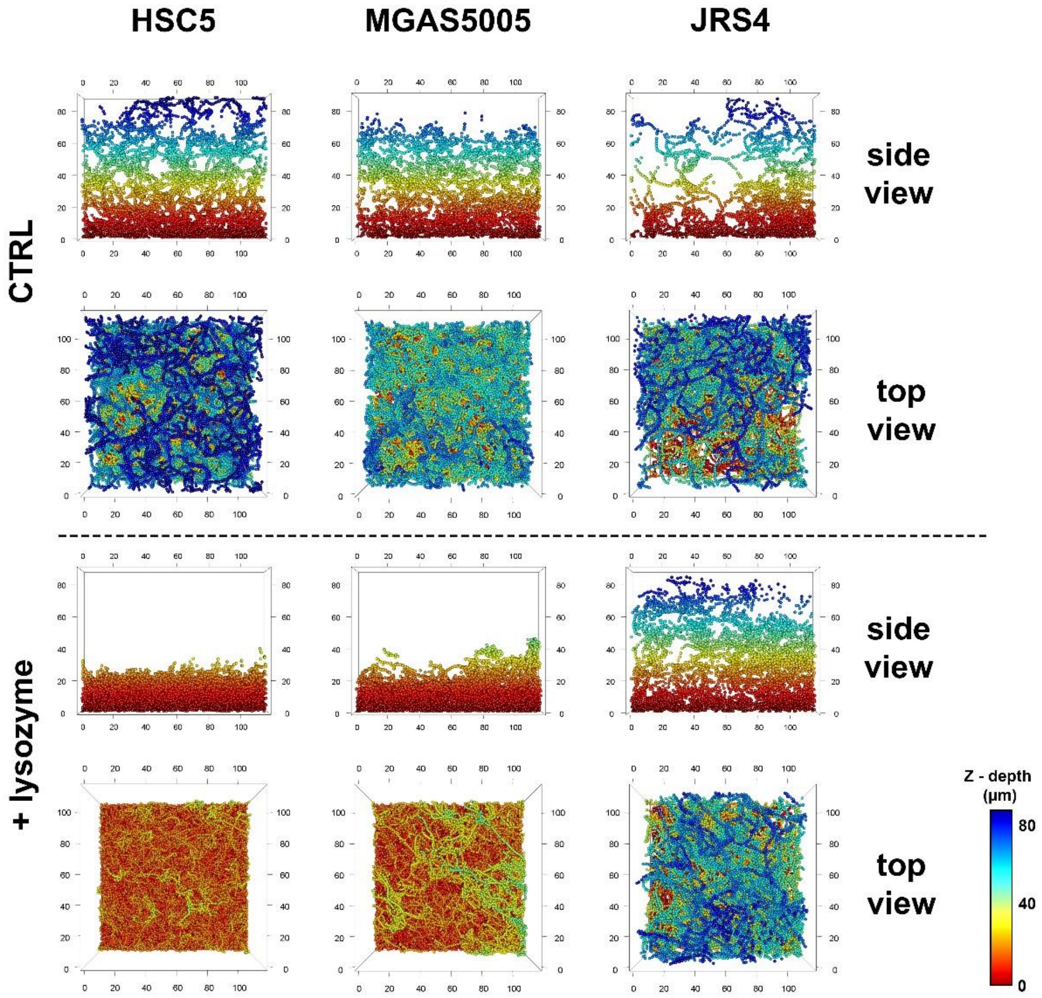


**Figure S3.** Structure visualization of HSC5, MGAS5005 and JRS4 biofilms grown without and with 10 µg/mL lysozyme (coloured by Z-depth). Field of view (XY): 130 × 130 µm (top view) and 130 x 10 µm (side view). Tick size: 20 µm.

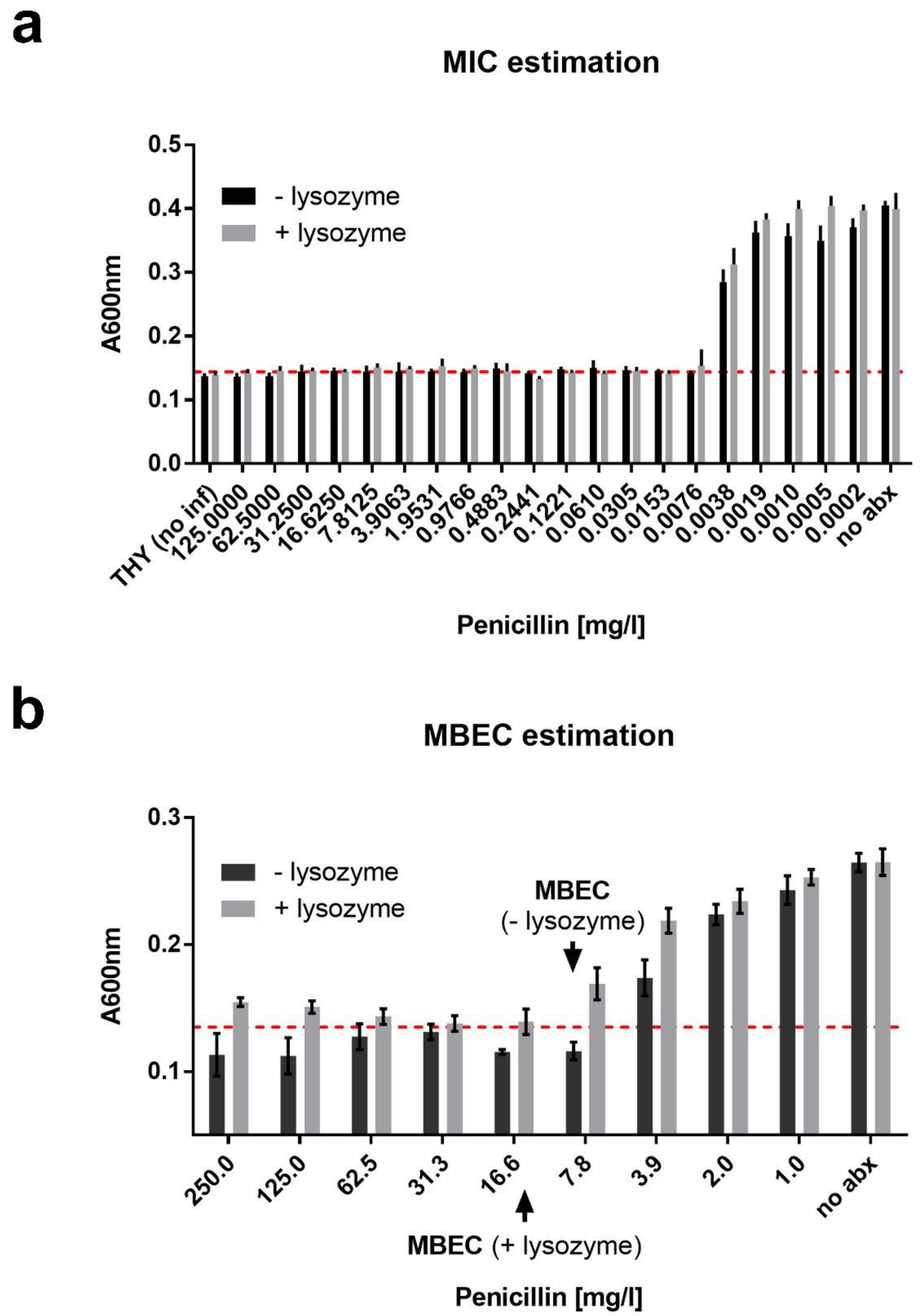


**Figure S4.** (a) MIC and (b) MBEC estimation for JS95 biofilms. Dashed line represents the average value from wells with no visible growth.

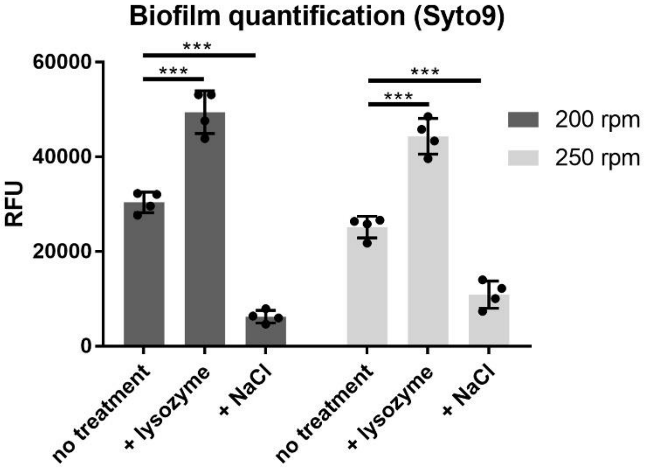


**Figure S5.** JS95 biofilm quantification using Syto9. Biofilms were grown with 10 µg/mL lysozyme or 0.9% total NaCl, as indicated in the figure. Dots and error bars represent biological replicates and mean ± SEM, respectively. Statistical analysis was performed using one-way ANOVA, followed by Dunnett's multiple comparisons test. ***, $P < 0.001$.

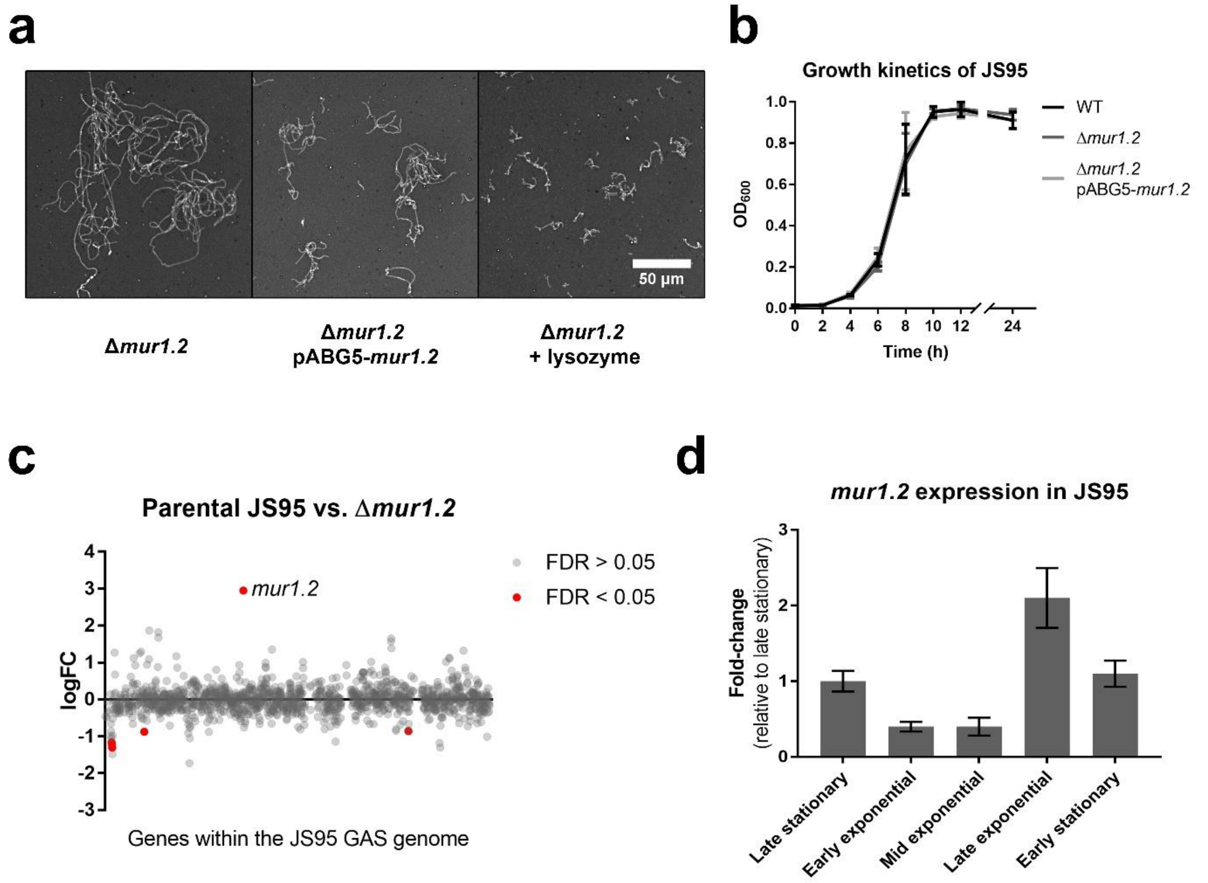


**Figure S6.** **(a)** Bacterial chaining from overnight cultures of the autolysin mutant (Δ*mur1.2*) and the complement strain (Δ*mur1.2* pABG5-*mur1.2*) in THY-G medium, compared to Δ*mur1.2* grown in THY-G medium supplemented with 5 µg/mL lysozyme. **(b)** Growth kinetics of JS95 WT, Δ*mur1.2* and the complement strain, when grown in THY-G medium. **(c)** RNA-seq between parent strain and Δ*mur1.2*. Initially identified differentially expressed genes that passed the threshold of FDR<0.05 (shown in red) were subsequently confirmed as false positives by qPCR (data not shown), except deleted *mur1.2*. **(d)** Gene expression levels of *mur1.2* across different growth phases. Expression of *mur1.2* was quantified by qPCR for JS95 strain grown in THY-G medium at early exponential ($OD_{600}$ = 0.2), mid exponential ($OD_{600}$ = 0.5), late exponential ($OD_{600}$ = 0.8), early stationary ($OD_{600}$ = 1.0), and late stationary (15 hpi) growth phases. Error bars represent mean ± SEM from two independent experiments.

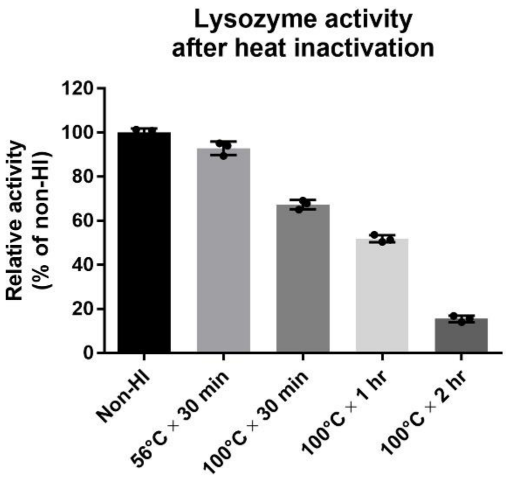


**Figure S7.** Chicken egg white lysozyme activity, as assessed by the EnzChek Lysozyme Assay Kit (Thermo-Fisher, USA), after heat inactivation with varying inactivation time and temperature. Error bars represent mean ± SEM.

### SUPPLEMENTARY METHODS

Table S1A. List of strains and plasmids used in this study.

| Strain or plasmid | Description | Reference |
|---|---|---|
| *S. pyogenes* JS95$_{ATG}$ WT | Derivative of NF isolate M14 JS95 containing a functional start codon in *silCR* | (1-3) |
| *S. pyogenes* JS95$_{ATG}$ Δ*mur1.2* | Deletion strain of *mur1.2* | This study |
| *S. pyogenes* JS95$_{ATG}$ Δ*mur1.2* pABG5-*mur1.2* | Complement strain of *mur1.2* | This study |
| *S. pyogenes* HSC5 | Wild-type strain | (4) |
| *S. pyogenes* MGAS5005 | Wild-type strain | (3, 5) |
| *S. pyogenes* JRS4 | Wild-type strain | (6) |
| *E. coli* DH5α pABG5 | *E. coli* DH5α with the pABG5 plasmid | (7) |
| *E. coli* DH5α pABG5-*mur1.2* | pABG5 containing *mur1.2* under the control of the *rofA* promoter | This study |
| *E. coli* DH5α pGCP213 | *E. coli* DH5α with the pGCP213 plasmid | (8) |
| *E. coli* DH5α pGCP213-Δ*mur1.2* | pGCP213 containing the flanking regions of *mur1.2* for deletion mutagenesis | This study |

Table S1B. List of primers used in this study.

| Name | Sequence (5’ to 3’) | Description |
|---|---|---|
| Generation of deletion construct | | |
| Mur12F1 | CGCGGATCCAATAATGGCAAGTCACAAC | Targets upstream of *mur1.2* |
| Mur12R1 | TTACATTAATAAGTCATGCGTGTCATTATTCACATATCCT | Targets upstream of *mur1.2* |
| Mur12F2 | AGGATATGTGAATAATGACACGCATGACTTATTAATGTAA | Targets downstream of *mur1.2* |
| Mur12R2 | CGGGGTACCCCAAATCTAAAACTGCTCAG | Targets downstream of *mur1.2* |
| M13-F | GTAAAACGACGGCCAG | Sequencing of insert |
| M13-R | CAGGAAACAGCTATGAC | Sequencing of insert |
| Generation of expression construct | | |
| ABG5X1F | ATCAGTCTGACGACCAAGAGAGC | To linearize pABG5 |
| ABG5X1R | TCAGTTCCTCACAATAATGGTTTAGTTGTTAAAAGG | To linearize pABG5 |
| Mur12X1F | ATTGTGAGGAACTGAATAAAAAAGGATATGTGAATAATGACAAAAAAGAAAGGTAAGC | Amplifies the *mur1.2* gene |
| Mur12X1R | GGTCGTCAGACTGATACAACGATAAACATATCAAAGTAAAACTAAAAGGATTTAGTTCT | Amplifies the *mur1.2* gene |
| ABG5F2 | TTGCCAATAACTGAGGTAGCA | For sequencing of insert |
| ABG5R2 | TTATGGCTCTCTTGGTCGTC | For sequencing of insert |
